# Mapping the soil microbiome functions shaping wetland methane emissions

**DOI:** 10.1101/2024.02.06.579222

**Authors:** Mikayla A. Borton, Angela M. Oliverio, Adrienne B. Narrowe, Jorge A. Villa, Christian Rinke, David W. Hoyt, Pengfei Liu, Bridget B. McGivern, Emily K. Bechtold, Jared B. Ellenbogen, Rebecca A. Daly, Garrett J. Smith, Jordan C. Angle, Rory M. Flynn, Andrew P. Freiburger, Katherine B. Louie, Brooke Stemple, Trent R. Northen, Christopher Henry, Christopher S. Miller, Timothy H. Morin, Gil Bohrer, Kelly C. Wrighton

## Abstract

Accounting for only 8% of Earth’s land coverage, freshwater wetlands remain the foremost contributor to global methane emissions. Yet the microorganisms and processes underlying methane emissions from wetland soils remain poorly understood. Over a five-year period, we surveyed the microbial membership and in situ methane measurements from over 700 samples in one of the most prolific methane-emitting wetlands in the United States. We constructed a catalog of 2,502 metagenome-assembled genomes (MAGs), with more than half of the 70 bacterial and archaeal phyla sampled containing novel lineages. Integration of these data with 133 soil metatranscriptomes provided a genome-resolved view of the biogeochemical specialization and versatility expressed over wetland soil spatial and temporal gradients.

Centimeter-scale depth differences best explained patterns of microbial community structure and transcribed functionalities, even more so than land coverage or temporal information. Moreover, while extended flooding restructured soil redox, this perturbation failed to reconfigure the transcriptional profiles of methane cycling microorganisms, contrasting with theoretical expected responses to hydrological perturbations. Co-expression analyses coupled to depth resolved methane measurements exposed the metabolisms and trophic structures most predictive of methane hotspots. Mapping the spatiotemporal transcriptional patterns on this compendium of biogeochemically classified soil derived genomes begins to untangle the microbial carbon, energy, and nutrient processing contributing to wetland methane production.

**Importance:** Soil microbial ecology is increasingly recognized as essential to climate mitigation, but realizing its full potential requires shifting from static genome inventories to dynamic assessments of microbial activity. This study shows that methane-cycling microbes exhibit stable, depth-stratified expression patterns, even in response to major redox and flooding shifts, undermining assumptions that water-table manipulations common in wetland management can alone reduce methanogenesis. Instead, methane cycling is shaped by spatially organized, transcriptionally active networks involving not only methanogens but also methanotrophs, fermenters, and iron reducers. These findings expose the limitations of genome-only models and highlight the need for soil diagnostics that capture in situ activity. Together, we provide a foundation for developing activity-based microbiome tools, embedding microbial functions into Earth system models, and designing interventions that move beyond “single lever” strategies and instead work with the structure and dynamics of microbial communities as complex, layered systems.

## Introduction

Wetlands are characterized as waterlogged soils, rich in organic matter, with low concentrations of oxygen in the porewater. Consequently, the anoxic microbial decomposition of this soil organic matter leads to the production of biogenic methane (CH_4_). As such, wetland soils are the largest and most variable source of this potent greenhouse gas, accounting for nearly a third of annual CH_4_ emissions (1–4). Despite this global climate relevance, the microbial communities and metabolic pathways driving the conversion of soil organic carbon to CH_4_, along with their environmental controls, remain unresolved. This knowledge gap hinders the incorporation of microbial processes into land surface and biogeochemical models, impeding the accurate management and predictions of greenhouse gas (GHG) fluxes from wetlands (3, 5, 6).

Over the last decade, metagenomic sequencing approaches have described the microbial community membership and function associated with CH_4_ production across a range of habitats. While other high CH_4_ emitting systems, like wastewater (7), ruminants (8), or subsurface (9), habitats have received considerable genomic scrutiny, wetland soils remain relatively underexplored in terms of microbial genomic sampling. Of the handful of genome-resolved studies from freshwater wetlands, nearly all are focused on organic soils from northern peatlands (10–13). One notable study, from thawing permafrost in Sweden, yielded ∼1,500 metagenome assembled genomes (MAGs) (11), serving as the stand alone microbial catalog for all wetland soils. This means the highest CH_4_-contributing wetland types, temperate and tropical marshes, have only a few genome-resolved studies to date (14–16). These studies emphasized the roles of a limited number of microorganisms to specific biogeochemical processes (14–17), such as methanotrophy, yet failed to inventory the overall composition, function, and interconnected metabolic pathways driving CH_4_ production across diverse wetland gradients.

To bridge this knowledge gap, we undertook a comprehensive sampling within a temperate freshwater marsh, aiming to elucidate the spatiotemporal related variations in the microbiome. This freshwater wetland was chosen due to its exceptionally high CH_4_ fluxes, exceeding ten times the median flux observed in wetlands with similar annual temperature ranges (18, 19), and because of its susceptibility to climate-induced flooding events influenced by fluctuating Lake Erie water levels (20). Leveraging more than 5 Tera basepairs of sequencing, we reconstructed thousands of microbial genomes and integrated this data with soil metatranscriptomes, geochemistry, and greenhouse gas porewater measurements and fluxes.

Here we delineate the expressed membership and metabolisms of the microbial community, shedding light on the biogeochemical contributions of hundreds of understudied lineages. These findings offer new perspectives on the stability and interactions of metabolic guilds driving CH_4_ production in temperate, freshwater wetland ecosystems. This study provides mechanistic insights that pave the way for activity-based methane mitigation strategies and more accurate integration of microbial processes into Earth system models (21).

## Results

### Sampling microorganisms from wetland soils

As a model wetland we selected the highest methane-emitting wetland in the AmeriFlux network (site ID US-OWC), an investigator network with tailored instrumentation, data processing, and standardized procedures for flux measurements (19, 22–24). The Old Woman Creek (OWC) wetland was instrumented to measure wetland CH_4_ at various spatiotemporal scales (**Supplementary** Fig. 1), facilitating a detailed assessment of the microbial processes that impact soil CH_4_ dynamics. Based on land coverage types, we parsed the wetland into 10 patches (**Fig. 1A**). These patches included 5 land coverage types including emergent vegetation-*Typha* (n=1), emergent vegetation-*Nelumbo* (n=3), standing freshwater or open water (n=3), and temporal mud flats (n=3). From selected sites, once per month during sampling campaigns, dialysis peepers measured soil CO_2_ and CH_4_ concentrations along ∼3cm-depth increments up to 30 cm deep. This content was paired to surface chamber flux measurements for CO_2_ and CH_4_ and site wide fluxes using the eddy covariance and meteorological station. Surrounding the peepers, soil cores were pulled and extruded at 5 cm depth increments down to 25 cm to result in 705 samples for soil geochemical, metabolite, and microbial analysis. In summary, across the five-year sampling campaign (2013-2018), each one of these 10 ecological patch sites was sampled at least 52 times for paired microbial, soil geochemistry, flux and porewater GHGs. The inventory of measured geochemistry and flux data and their relationships is shown in **Supplementary Fig. S2**.

**Figure 1.**
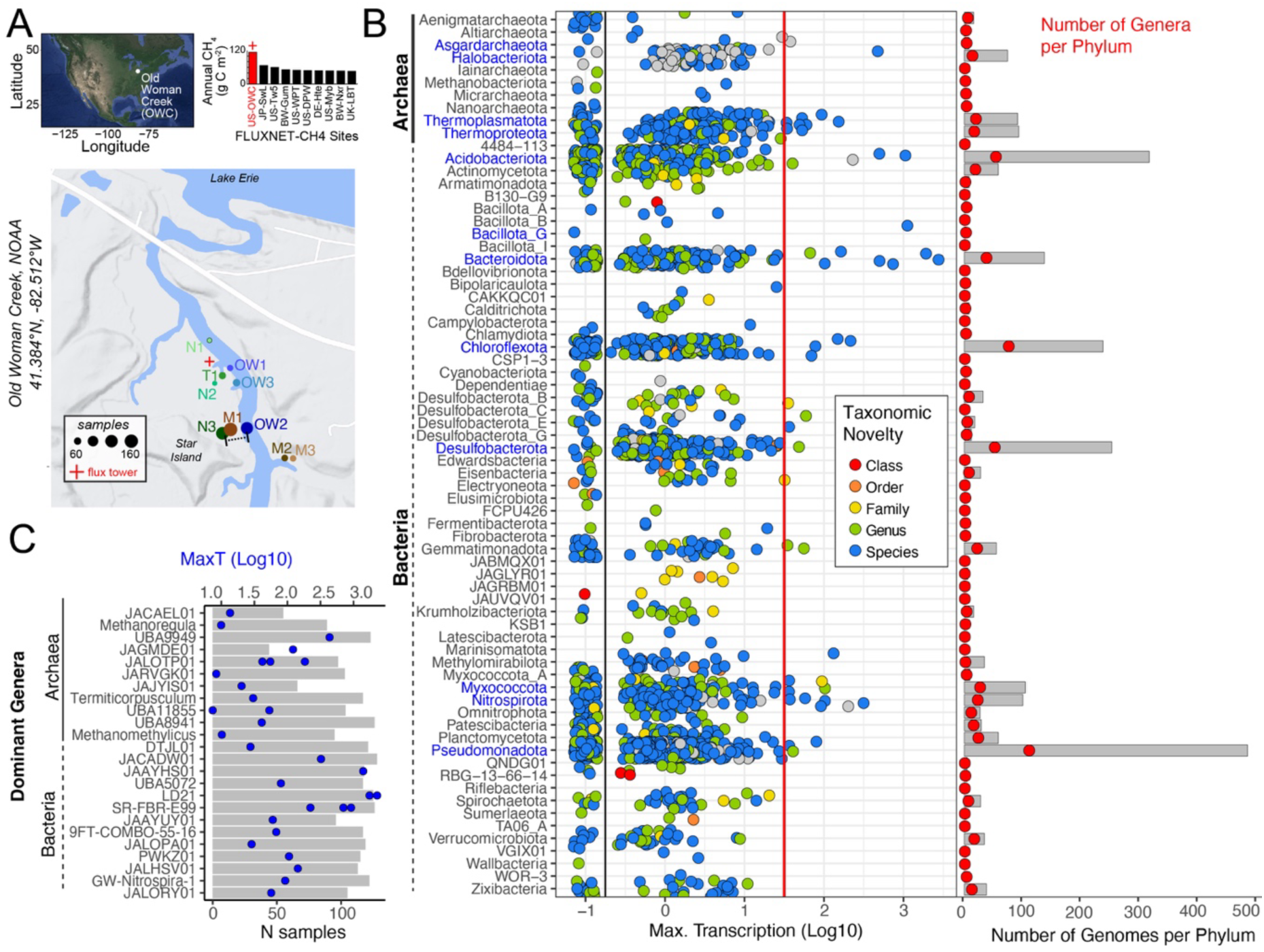
Freshwater wetland soils host thousands of taxonomically diverse MAGs that recruit metatranscripts. **A)** Old Woman Creek (OWC) is located on the southern part of Lake Erie and is the highest US methane emitting wetland in the AmeriFlux Network. A site map of the OWC freshwater wetland shows where the 700 samples were collected from 10 major sampling sites that were historically defined as emergent vegetation, either *Nelumbo* (N1-3) or *Typha* (T1), mud cover (M1-3), or location in open water channel (OW1-3). Sites were sampled over multiple years and depths at 5 cm intervals from 0-30 cm, with sampling density proportional to circle size, 52 is the minimum sampling per location (**Supplementary Fig. S1**). **(B)** The taxonomic distribution of the 2,502 MAGs summarized at the phylum level. Bold phylum label indicates a candidate phylum, and blue text indicates the phylum contains a top 12 dominant genus (shown in **1C**). Each point represents a MAG and is colored by taxonomic novelty. Points are distributed by their log10 maximum transcription across the 133 metatranscriptome samples. MAGs to the right of the black line were considered positive for sufficient transcript recruitment (N = 1,948). MAGs to the right of the red line had the greatest transcription (log10 max. transcription > 1.5). The gray bar chart to the right summarizes the number of genomes per phylum recovered, and red points overlaid represent the number of genera recovered within each phylum, using the same x-axis. **(C)** The transcriptionally most dominant archaeal and bacterial MAGs (top 12 for each domain) from 1B, grouped by genus. Gray bar indicates metatranscriptome prevalence (gray bar), or the number (N) of the 133 samples where metatranscripts were recruited to the MAG for that genus. The axis is shown on bottom (N samples). The blue dot indicates the maximum transcription from **1B**, with the axis (MaxT, Log10) shown at the top of the graph.

We screened 671 of these collected soil samples with 16S rRNA amplicon sequencing for methanogen membership (**Supplementary Table S2**), identifying 42 samples for deep metagenomic sequencing (**Supplementary Table S1**). We generated up to 300 Gbp of sequencing per sample, resulting in 3.1 Tbp total metagenomic sequencing (**Supplementary Table S3**). Collectively, this effort doubled the amount of metagenomic sequencing from wetland soils available in public repositories (**Supplementary** Fig. 1D). Given that one of our primary objectives was to increase the recovery of methanogen genomes from soils, multiple assembly and binning methods targeted the recovery of these lineages (**Supplementary** Fig. 1E). This approach resulted in over 17,333 draft genomes, which were dereplicated into a final collection of 3,217 genomic representatives, with 2,502 of these constituting high and medium quality metagenome assembled genomes (MAGs) (**Supplementary Fig. S1**).

To map which microbial genomes recruited transcripts and which metabolisms were expressed across sites, depths, seasons, and years, we obtained 2.7 Tbp of metatranscriptome sequencing from 133 soil samples. Nearly 80% of the metatranscriptomes had paired NMR detected metabolites to profile methanogenic substrates across the sample set (**Supplementary Table S4**). This comprehensive multi-omics sampling across wetland gradients, including multiple land coverage types, depths, and times, provides the first insights into the microbial structural and functional diversity residing in temperate freshwater wetland soils.

### Uncovering thousands of transcribed genomes from wetland soils

We reconstructed MAGs from 60 bacterial (N=2,204 MAGs) and 10 archaeal (n=298 MAGs) phyla that included 85 methanogen-related MAG representatives. Many MAGs recovered here were designated newly sampled lineages at the genus level or higher including genome representatives with unnamed classes (n=4 MAGs), orders (n=20 MAGs), families (n=15 MAGs), and genera (n=122 MAGs) (**Supplementary Table S3**). In addition, hundreds of genomes were from poorly or previously uncharacterized phyla often denoted only by an alphanumeric identifier or placeholder name, such as QNDG01, CSP1-3, and KSB1 (**Fig. 1B**). In this wetland, the most represented phyla are the archaeal Halobacteriota (n=74 MAGs) and the bacterial Pseudomonadota (n=484, formerly Proteobacteria), Acidobacteriota (n=316 MAGs), Desulfobacterota (n=252), and Chloroflexota (n=237 MAGs). The reconstruction of these thousands of genomes, many metabolically uncharacterized, highlights the vast, unsampled microbial diversity still residing within soils.

Approximately 75% of the reconstructed genomes (n=1948) recruited transcripts from the 133 metatranscriptomes. We report the maximum transcription for each MAG and denote the top 12 most transcriptionally dominant MAGs from each bacterial or archaeal domain (**Fig. 1BC**). These MAGs with dominant transcription also have high transcriptional occupancy, with a minimum detection in 35% across the 133 metatranscriptome samples (**Fig. 1C**). A significant majority (83%) of these transcriptionally dominant genomes belong to poorly characterized genera as noted by their names (e.g., genera defined only by alphanumeric code). The prevalence of transcriptionally dominant but poorly classified genomes reveals a gap in our understanding of the organisms actively shaping wetland biogeochemistry.

### Connecting transcribed biogeochemical functions to microbial taxonomy

Each of the 1,948 MAGs that recruited transcripts was assessed for expressed gene content relating to 14 biogeochemical processes: oxygen (aerobic respiration, microaerophilic respiration), methane (methanogenesis, aerobic methanotrophy, anaerobic methanotrophy), sulfur (sulfur oxidization, sulfur reduction), nitrogen (nitrification, nitrogen reducer, nitrogen fixation, and Dissimilatory Nitrate Reduction to Ammonium (DNRA)), iron (iron oxidation, iron reduction), and other metabolisms (phototrophy, obligate fermentation) (**Supplementary Fig. S3, Methods**). This resulted in 1,135 MAGs that contained sufficient gene transcript recruitment to be assigned to at least one of these 14 biogeochemical guilds (**Supplementary Table S5)**.

Aerobic and microaerophilic metabolisms were the most widely encoded across wetland lineages (**Fig. 2A**). We compared the dominant encoded metabolic traits (i.e., detected in the greatest number of genomes) to the most dominant transcribed traits, finding the transcribed functionalities did not follow the encoded (**Fig. 2B**). For example, the capacity for microaerophilic and aerobic respiration was detected in the largest percentage of genomes (53% and 51%) but only detected in 8% and 6% of genomes in our transcriptomes respectively. In contrast, transcription for genes enabling anaerobic metabolisms like obligate fermentation, iron reduction, and nitrogen reduction were detected more so than aerobic metabolisms. This enrichment for anaerobic metabolisms in the transcript signal is not entirely surprising given that most soils in our data set have pore-water dissolved oxygen (DO) concentrations below the 1uM detection limit (**Supplementary Table S4**). Our approach reveals that transcriptional profiles capture environmentally responsive metabolic activity not evident from genome content alone, underscoring the importance of expression-based profiling to uncover active microbial processes.

**Figure 2.**
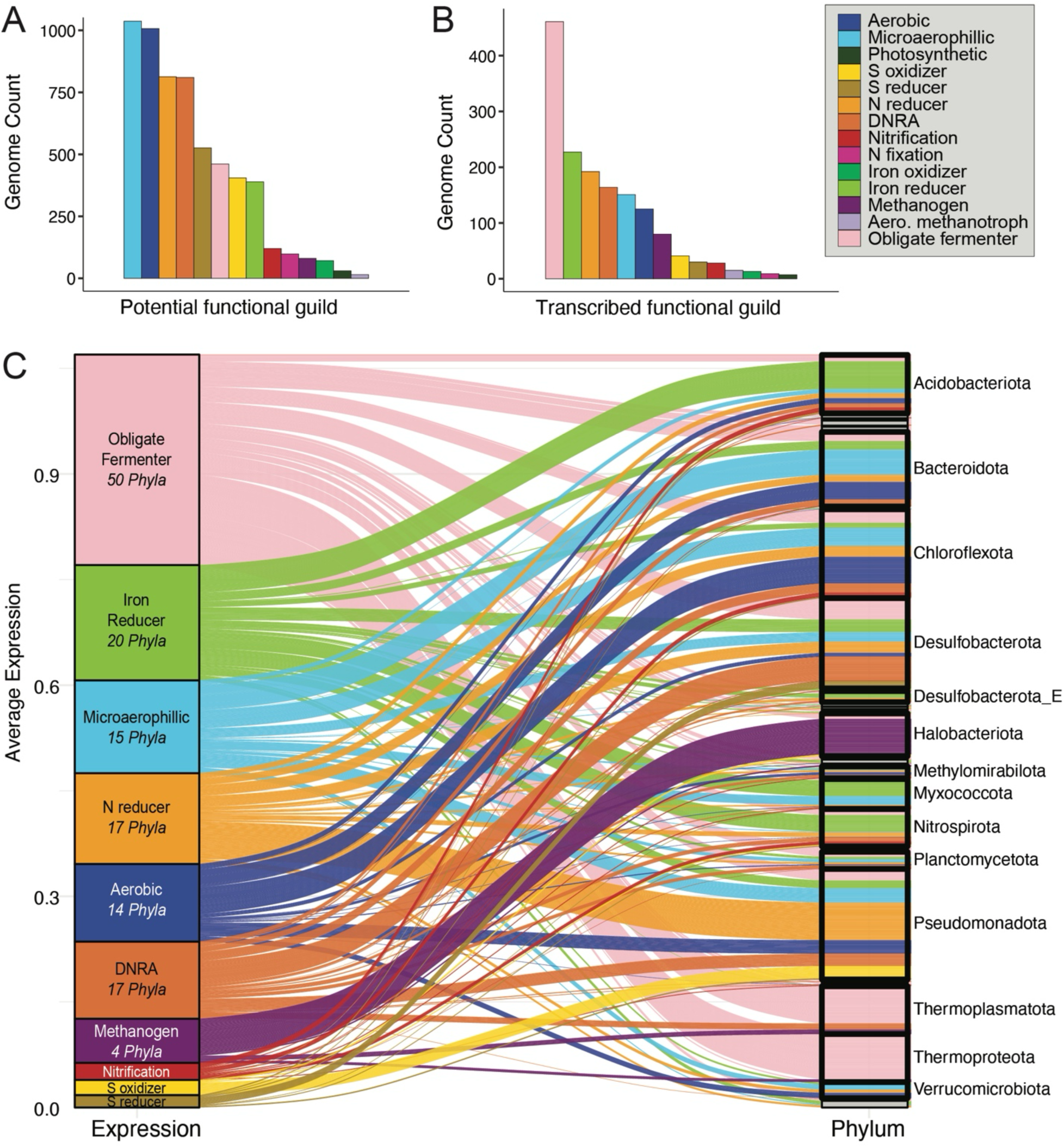
Distinct functional potential and expression across wetland microbial genomes. Bar plots summarize the genome counts across functional traits for genomic potential **(A)** and transcription **(B)**, respectively. **(C)** Alluvial plot illustrates the distribution of top 10 expressed metabolic guilds (left) and their corresponding phyla (right). The height of each flow represents the average expression of that metabolic guild in a particular phylum, with colors indicating different metabolic guilds. Labels for phyla with less than 0.01 total expression were omitted. Number of phyla for each guild is noted for 7 of top 10 phyla, with Nitrification (n=11 Phyla), S oxidizer (n=4 Phyla), and S reducer (n=7 Phyla) guilds detailed here.

Focusing on transcribed functions, at the phylum level we found that certain phyla, such as Thermoproteota or Halobacteriota, are characterized by a pink or purple color, indicating their specialization as obligate fermenters or methanogens (**Fig. 2C**). In contrast, phyla like the Pseudomonadota and Acidobacteriota exhibit a wide array of colors, signifying their trait multifunctionality at the phylum level. This analysis also reveals disproportional contributions from phyla to nutrient and redox transformations. For instance, MAGs transcribing genes for sulfur reduction are enriched in the Desulfobacterota, iron reduction are prevalent in the Acidobacteriota and Nitrospirota, aerobic respiration are prevalent across the Chloroflexota, and nitrogen reduction and sulfur oxidation are enriched in some lineages of the Pseudomonadota.

Notably, keystone processes like nitrification and nitrogen fixation were both rare (<5% of MAGs) and phylogenetically dispersed (**Supplementary Table S5**). Nitrogen fixation, for example, was detected in transcripts from just nine MAGs across three phyla that encompassed aerobes, iron reducers, and denitrifiers, as well as a methanogen and a nitrifier, illustrating how assimilatory and dissimilatory processes are coupled. This trait mapping reveals the phylogenetic distribution of key metabolic activities and underscores the diverse, active roles that microorganisms play in driving nutrient and redox transformations.

### Identifying the dominant drivers shaping wetland microbiome structure and function

The relative expression of genomes across depth profiles revealed distinct microbial community structures in the Plant, Mud, and Open Water habitats (**Figure 3**). In Plant habitats, Pseudomonadota dominated the surface layers (0–5 cm), while deeper layers (15–25 cm) showed an increased abundance of Desulfobacterota and Bacteroidota; similar phyla were present in Mud habitats, but Desulfobacterota was more consistently abundant throughout the profile, with Verrucomicrobiota becoming more pronounced at depth. In Open Water habitats, Pseudomonadota were again prominent near the surface, but deeper layers shifted towards Gemmatimonadota and Thermoplasmatota, suggesting increased anaerobic processes with depth. These patterns reflect habitat differences in microbial activity and adaptation along the redox gradient within each environment.

**Figure 3.**
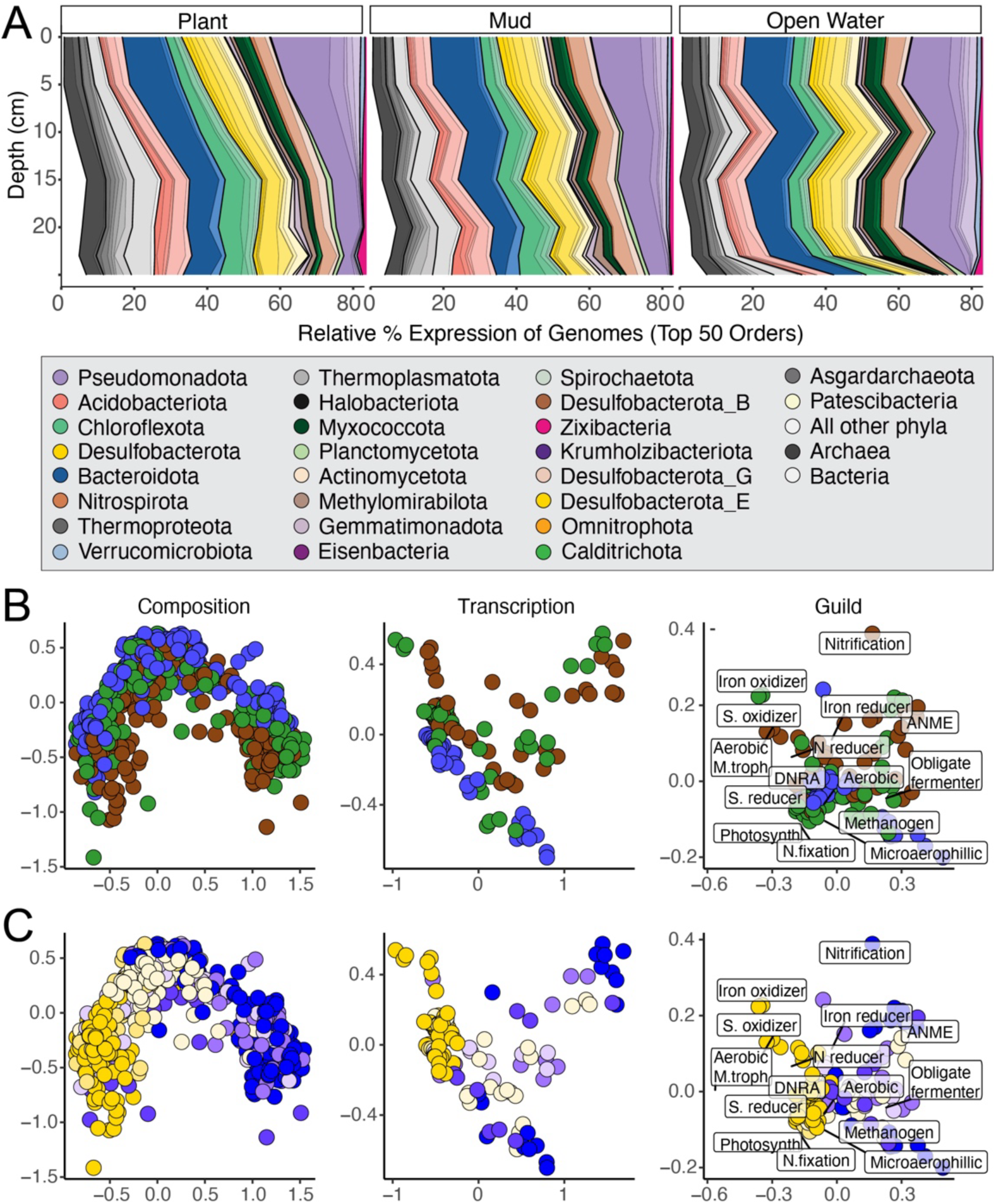
Depth structures wetland soil microbiomes. **A)** Smoothed density plots for the top 50 orders, shaded by phylum level colors provided in the legend. Changes in the relative expression across the sampled depth gradient and across land coverage types (vegetative, mud, and open water). **(B-C)** Nonmetric multidimensional scaling (NMDS) ordinations of wetland microbiome structural metrics are organized from (left) overall composition (N=671), (middle) genome transcription (N=133), and (right) guild transcription (N=133).The samples are colored by land coverage in (**B**) (brown=mud; green=vegetative; blue=open water), while **(C)** is colored by depth (yellow=surface to blue=deep). B and C highlight the responses of the two major environmental predictors with depth being the strongest predictor across all soil microbiome attributes (**Supplementary Figure S4**).

To test how different wetland environments influenced the soil microbiome, we evaluated the impact of 17 spatial, temporal, and geochemical factors across three levels of biological organization: (i) phylotype membership, (ii) transcriptionally ‘active’ genome membership, and (iii) expressed functional guilds (**Fig. 3BC**). Our analyses revealed that, regardless of the microbiome attribute measured, depth and depth-related geochemistry (such as soil Fe (II) and cation exchange capacity, CEC) were the best predictors of wetland soil microbiome structure and function, more so than differences from the different land coverages. These chemical and physical soil attributes played a more direct role in shaping expressed functional guilds, explaining 40 to 50% of this variation, nearly double variation explained for the phylotype data (**Supplementary Fig. S4**). Overall, this information can inform wetland microbiome sampling priorities, showing the value of adding more depth-resolved measurements and affirming collection of high-priority geochemical variables, such as CEC, Fe (II), pH, and acetate, while also recognizing that porewater oxygen measurements using hand held probes may fail to resolve redox microsite heterogeneity that drives wetland biogeochemical activity (15).

### Uncovering newly defined archaeal roles in carbon cycling

A valuable component of our sampling was the broad recovery of archaeal genomes (n=298 MAGs), spanning 10 phyla with 48 families including 88 genera. When considering the functional roles of Archaea in wetlands, these microorganisms were historically regarded primarily as methanogens or metabolizing other single carbon (C1) compounds (16, 25–27). Here we derived a gene-based rule set to assign these genomes to carbon decomposition guilds of (i) polymer hydrolysis, (ii) sugar oxidation, (iii) organic acids and nitrogen conversions, and (iii) C1 metabolisms (**Supplementary Table S7**).

We show phylogenetically diverse archaeal lineages expressed genes for hydrolyzing these plant polymeric constituents (**Fig. 4**). Depolymerization results in the second trophic group, yielding sugars and other oligomers or monomers. Lineages like Thorarchaeaceae and UBA233 transcribed genes to use 6 or more substrates that span both polymeric and sugar trophic groups. We also identified sugar utilizing specialist archaea (e.g., members of the TCS64) that expressed genes for utilizing at least 3 sugar substrates. In support of these expressed sugar metabolisms, raffinose, sucrose, and maltose like compounds were prevalent across the metabolome data (**Supplementary Table S8-S9**), giving further credence to the likelihood of these metabolisms in situ. These findings provide evidence that Archaea, often overlooked in primary carbon degradation, contribute far more broadly to soil carbon turnover than previously recognized, occupying roles across multiple carbon trophic levels.

**Figure 4.**
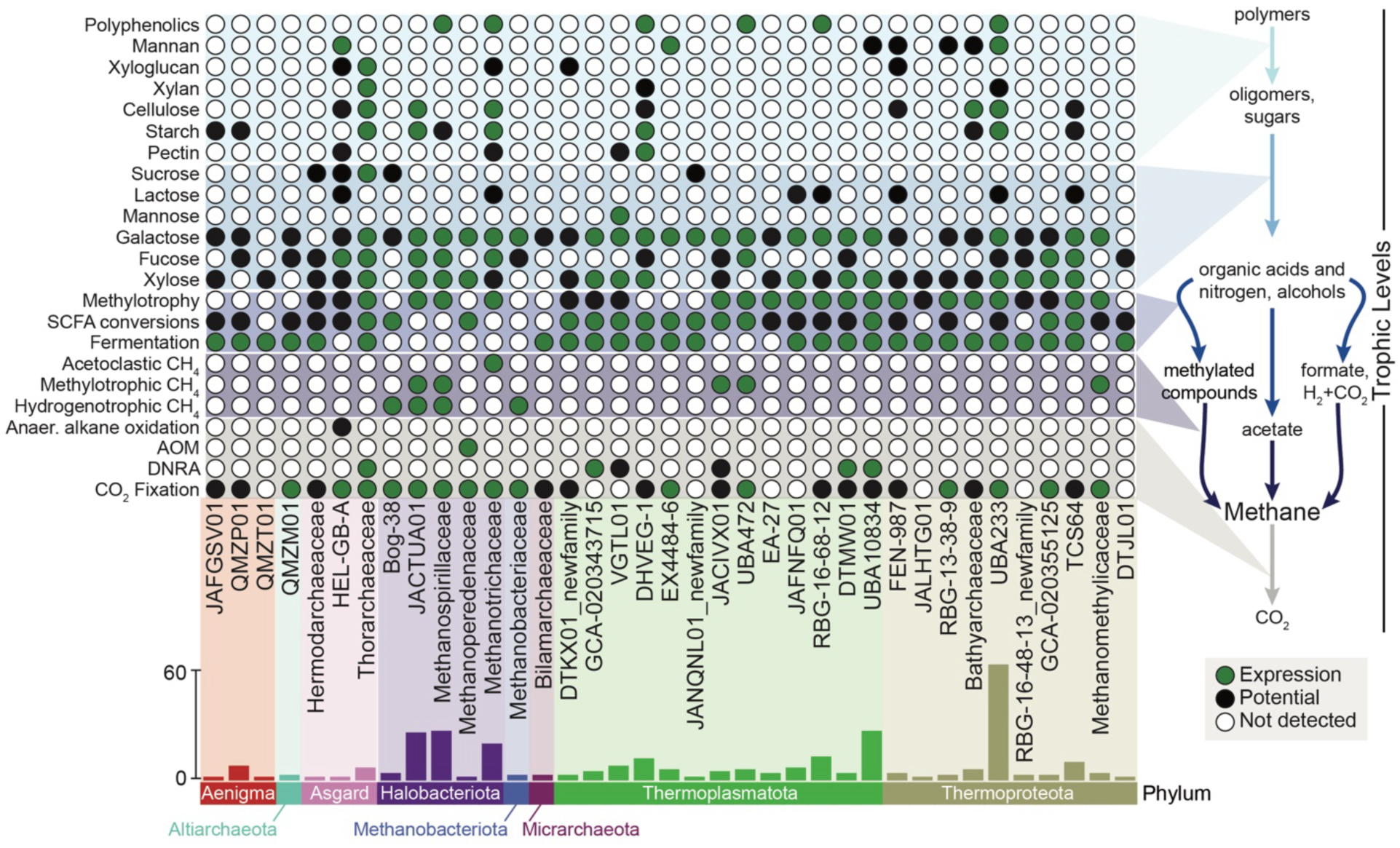
Assigning Archaeal roles in the wetland carbon cycle. (**Right**) The schematic on the right visualizes the major trophic levels contributing to soil carbon decomposition including: light blue polymers (polyphenolics, mannan, xyloglucan, xylan, cellulose, starch, pectin); medium blue oligomers and sugars (sucrose, lactose, mannose, galactose, fucose, xylose); dark blue organic acids, nitrogen, and alcohols (fermentation, methylotrophy, anaerobic alkane oxidation); purple methanogenesis (acetoclastic, methylotrophic, and hydrogenotrophic); and grey other 1 carbon metabolisms (anaerobic methane oxidation, CO_2_ fixation). (**Left**) For each archaeal family the bubble plot denotes genes transcribed (green), genes present (black), or genes absent (white) for the major carbon utilization pathways assigned to the trophic levels. (**Bottom**) The bar chart summarizes the number of transcriptionally active MAGs recovered within each archaeal phylum and family.

The third guild before methane production yields small molecular weight compounds like organic acids (e.g., acetate, butyrate, lactate) and methylated (e.g., methylamine, methanol) compounds. Some of these are methanogenic substrates, while others can be further fermented, typically syntrophically, to yield methanogenic substrates. This latter guild had the greatest transcriptional prevalence across the archaeal genomes, indicating the critical role these archaea, most of them diagnosed as obligate fermenters, could play in cross feeding methanogens.

The fourth guild of carbon is C1 metabolisms, with nearly a third of the archaeal genomes assigned as methanogens, representing 4 phyla (Methanobacteriota, Thermoproteota, Halobacteriota, Thermoplasmatota). The 85 methanogen MAGs spanned 10 families with 20 genera, 8 of the latter were new genera first identified here (**Supplementary Fig. S5**). Beyond canonical methanogens, we also recovered genomes from presumed non-methanogenic, *mcrA-* containing taxa such as the anaerobic methanotrophic *Methanoperedens* genus and the alkane-oxidizing Helarchaeales. Prior to this effort, the largest wetland MAG database contained 21 methanogen representatives with 4-fold less genus level diversity (11). Despite their direct climate feedback role, it is astonishing that multiple new, highly active methanogens can still be recovered from a single wetland.

Beyond taxonomic novelty, metabolic reconstruction with paired transcriptional profiles offered new insights into wetland methanogenesis (**Supplementary Fig. S5**). Acetoclastic methanogenesis was confined exclusively to *Methanothrix* and closely related members (16 MAGs). Our field data reinforce laboratory studies of *Methanothrix* and *Methanosarcina* (28), indicating that obligate acetoclastic *Methanothrix* outcompetes the facultative acetoclast *Methanosarcina* in situ due to more efficient substrate acquisition under the low acetate concentrations typical of these wetland soils (mean 45 µM; **Supplementary Fig. S6**). Across the methanogens, obligate hydrogenotrophy was the most prevalently transcribed pathway in 52% of the MAGs, with two of the most transcriptionally active Archaea identified as obligate hydrogenotrophs (*Methanoregula* and UBA9949, **Fig.1**). This work confirms and extends canonical methanogenic paradigms by showing not just potential but active dominance of *Methanothrix* and *Methanoregula* under natural, low-substrate field conditions.

While wetland CH_4_ is commonly attributed to acetoclastic and hydrogenotrophic methanogens (3), our findings highlight the overlooked importance of methylotrophic pathways. We found that 25% of wetland methanogen MAGs expressed genes for methylotrophic methanogenesis and one of the most transcriptionally active Archaea, *Methanomethylicus*, was a methylotroph (**Fig. 1C**). Additionally, we detected methylotrophic gene expression in lineages traditionally considered hydrogenotrophic, including *Methanoregula* MAG and in one MAG belonging to a novel genus within the *Methanospirillaceae* family (**Supplementary Fig. S5**). We further resolved substrate preferences for methylotrophic methanogens, showing specificity for methylated nitrogen, sulfur, and oxygen compounds. Supporting this, NMR and LC-MS analyses frequently detected methyl-N compounds (e.g., betaine, carnitine) and methyl-O compounds (e.g., methanol, vanillic acid) at higher concentrations and in more samples than canonical substrates like formate or acetate (**Supplementary Fig. S6**). These findings reinforce an emerging consensus that methylotrophic methanogenesis may be a widespread and underappreciated source of methane in wetland ecosystems (16, 27, 29).

### Methane cycling guilds are temporally stable along a depth gradient over years

Methane cycling genome-based transcription patterns from a flooded mud-flat collected monthly over a 3-month summer 2018 season exposed methanogen and methanotroph spatiotemporal niches and possible depth-defined ecotypes (**Supplementary Fig. S7**). We uncovered six core methanogens, from diverse functional and phylogenetic backgrounds, with high levels of mean transcription across all soil depths and timepoints. The remaining CH_4_ cycling members were localized to specific soil depths, forming co-expressing clades that persisted across this summer season. We report a greater proportional enrichment of acetoclastic and aerobic methanotrophs in surface soils, while the middle and deepest soil layers included the anaerobic methane-oxidizing *Methanoperedens* genome and proportionally more hydrogenotrophic and methyl-utilizing methanogens. Reflecting these latter methanogens requirements for hydrogen, we observed increased formate concentrations from deeper (10-30 cm) depths across our mud, plant, and open water land coverage patch types (**Supplementary Fig. S6**). Our findings highlight the persistence of depth-stratified methane ‘neighborhoods’, composed of many closely related strains with a high-degree of functional redundancy.

Given the seasonal stability across summer 2018, we next compared the multi-omics data collected from this site in August 2018 to parallel samples from the same mud flat in August 2015. During this time, the wetland flooded and the exposed mudflat from 2015 was beneath 5.3 ft of overlying freshwater in 2018. Based on existing theory about the redox sensitivity of methanogens and their responses to hydrological perturbations (30), with flooding we hypothesized to observe decreased dissolved oxygen (DO) concentrations and increased methanogen expression broadly across the community, particularly in surface soils. Matching flooding expectations, DO decreased to non-detectable levels in surface soils subsequent to flooding. In contrast to expectations, 90% of the methanogen and aerobic methanotroph (Methylococcaceae, Methylomonadaceae) MAGs experienced no significant change in mean genome transcription in the flooding restructured surface soils (**Fig. 5**). Worth noting, a few taxa were flooding-responsive, like the methylotrophic *Methanomethylicus* (**Fig. 1C**). Supporting this lack of restructuring in the CH_4_ cycling microbiome, the concentrations of methanogenic substrates like formate, methanol, and acetate did not change with flooding (**Supplementary Fig. S6**). This flooding response may be due to the fact that methanogens in these routinely submerged and drained soils surface soils could be already sheltered within anoxic microsites, such as biofilms or soil aggregates (15, 31).

**Figure 5.**
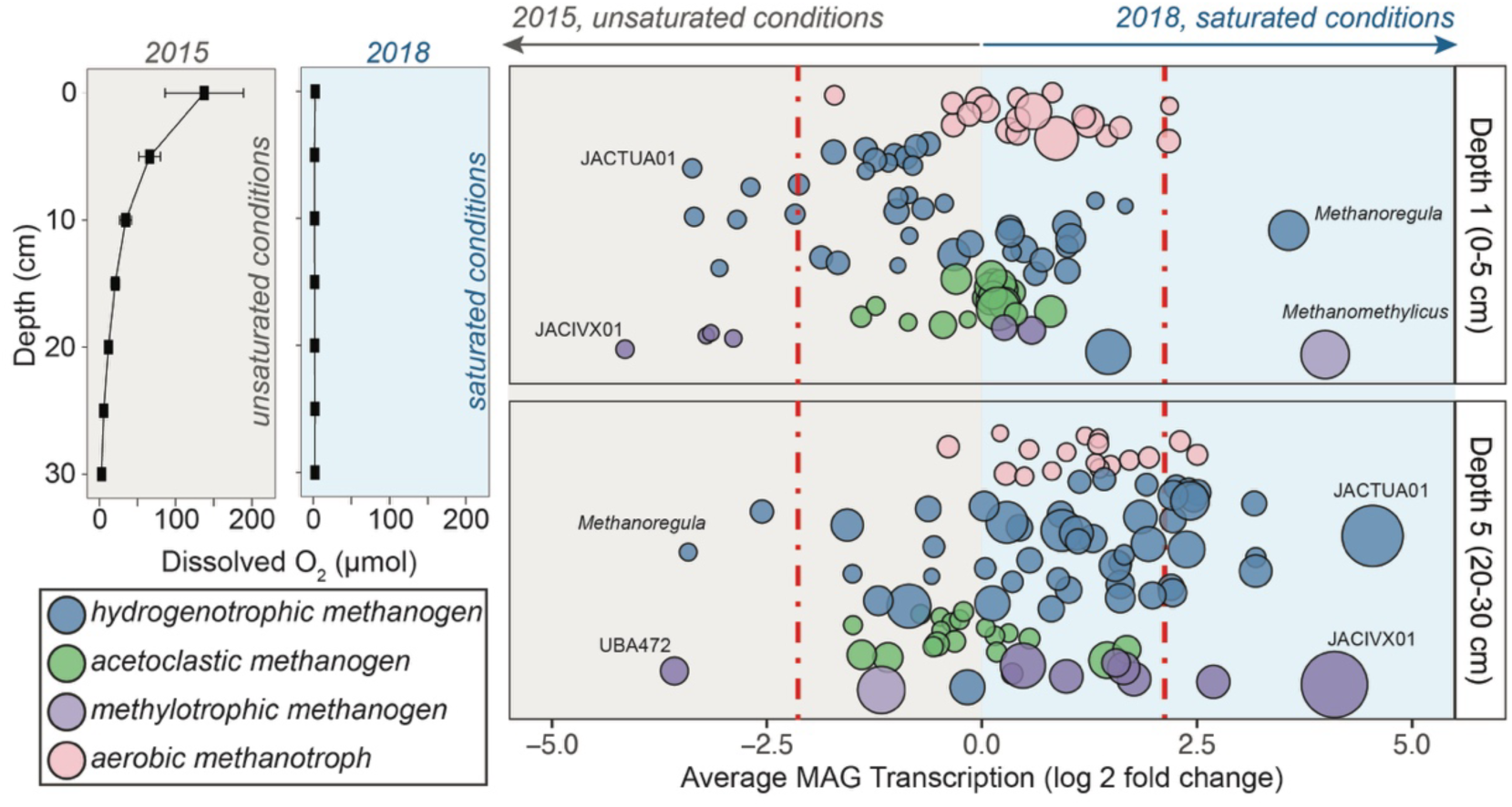
Methanogen stability in response to flooding. Dissolved porewater oxygen depth profiles from August 2015 (grey) and 2018 (blue) highlight the redox impact of flooding. Scatterplot shows the change of mean MAG transcription from 2015 to 2018 from the methane cycling community within surface (0-5 centimeters) and deep (20-30 centimeters) soil compartments. Bubbles represent a single methane cycling MAG and are colored by metabolism and sized by mean transcription. MAGs within the red dashed lines do not have log2 fold change in mean transcription and are considered temporally stable.

### Co-expression networks reveal key lineages modulating soil greenhouse gasses

We implemented a workflow to integrate the data collected across this manuscript to render the carbon and energy trophic network predictive of soil CH_4_ concentrations (**Supplementary Fig. S8**). First, weighted gene co-expression identified 7 modules composed of genomes with shared transcriptional patterns across our 133 metatranscriptome samples (**Fig. 6A**). Two of these modules predicted soil porewater CH_4_ concentrations, with the surface-soil associated turquoise module negatively correlated and the deeper-soil associated brown module positively correlated to soil CH_4_ concentrations (**Fig. 6A**). These surface and deep modules contained 551 and 372 MAGs, respectively. The top ten microbial genomes with the greatest importance for CH_4_ prediction and their transcribed biogeochemical traits are also noted (**Fig. 6CD**). Distilling the transcribed carbon and energy functional gene content of these predictive genera, we mapped the trophic configurations that contribute to CH_4_ concentrations in these depth-defined compartments (**Fig. 6E**).

**Figure 6.**
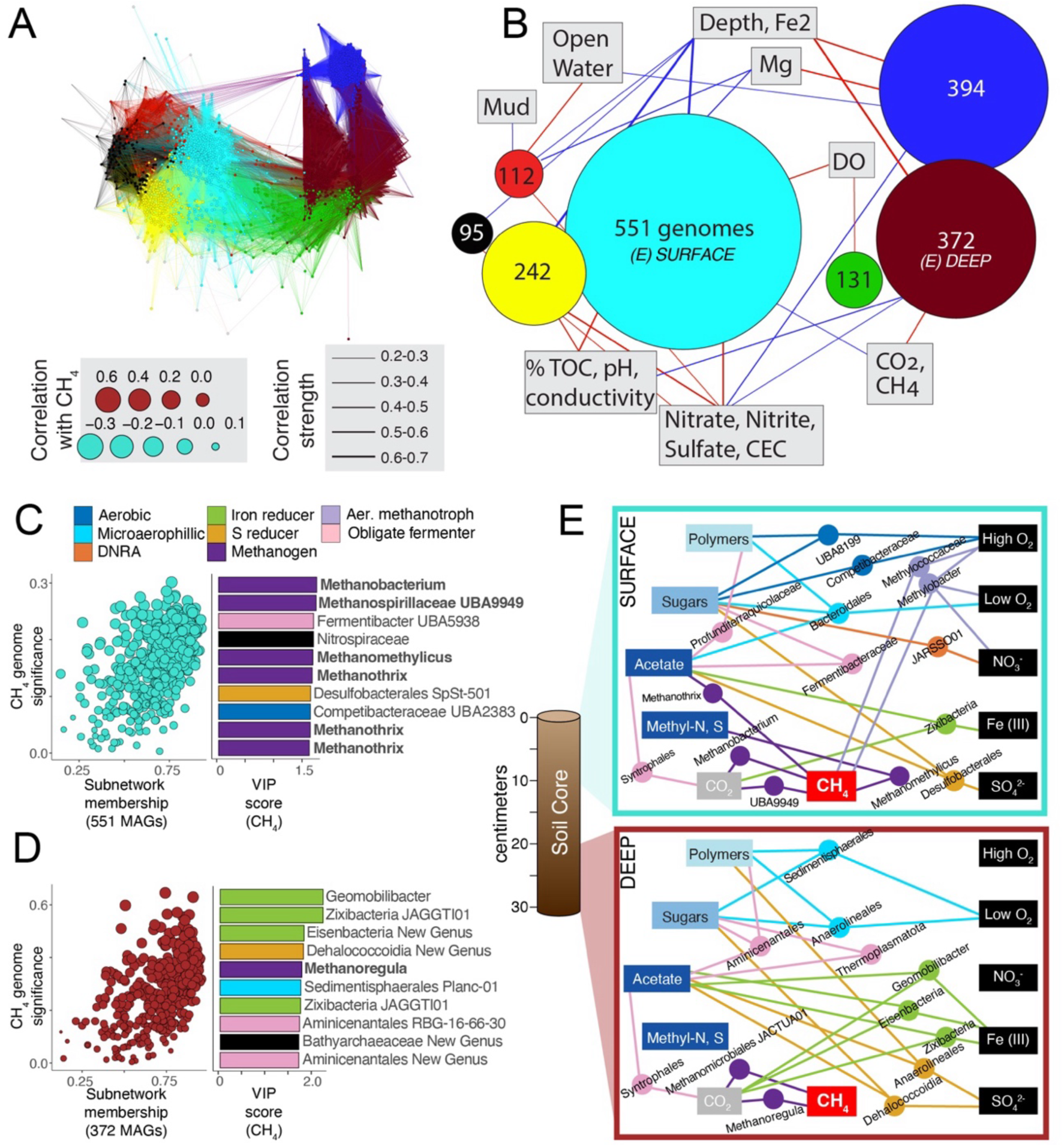
Coordinated gene expression networks predict wetland methane concentrations. **(A)** Overall co-expression network from 1,948 MAGs across 132 metatranscriptomics samples. Node outline colors highlight modules (where N > 100) identified with weighted gene co-expression analyses. **(B)** A simplified network diagram highlighting the associations between modules (sized scaled by number of MAGs) and environmental features. Red and blue edges correspond to positive and negative associations, respectively. **(C)** The genome significance plot and highest variable important genomes related to CH_4_ concentrations in the turquoise, surface derived subnetwork (n = 551 MAGs, Rho = 0.53, P < 0.001). **(D)** The genome significance plot and highest variable important genomes related to CH_4_ concentrations from the brown, deep subnetwork (n = 372 MAGs, Rho = 0.58, P < 0.001). For panels C and D, the bar charts (middle) highlight the relative contribution of the top 10 variably important genomes (colored by metabolic classification). Methanogens are highlighted with bold text. **(E)** A process-based schematic based on genome-resolved transcript data of the significant variable important projection (VIP, >1) taxa and their contribution to methane production in the surface subnetwork from A (top) and the deep subnetwork from B (bottom).

Consistent with our taxonomic-based analyses, we found key biogeochemical processes (such as iron reduction, microaerophilic respiration, and methane production) were transcribed across depths in these predictive modules (**Fig. 6CD**). The iron-reducing Zikibacteria and the obligate fermenter Syntrophales were important predictors in both depth compartments However, distinct microbial lineages were often responsible for mediating the same functions at different depths. For instance, microaerophilic respiration involved *Bacteroidales* in surface soils, while *Anaerolineales* and *Sedimentisphaerales* fulfilled similar roles in deeper soils. The turquoise surface module also included functions not observed in the deep module, such as nitrate reduction and low-affinity cytochrome oxidase expression, traits adapted to higher oxygen conditions. Additionally, transcriptional evidence for aerobic methanotrophy (*Methylococcaceae* UBA6136 and *Methylomonadaceae* KS41) was exclusive to the surface module. This likely explains the negative correlation between the turquoise module and CH₄ concentrations in surface soils (**Fig. 6B**).

Together, these transcriptome-derived, process-based observations reveal expressed biogeochemical interactions in wetland soils. Soil CH₄ concentrations appear to be emergent properties of the microbiome, driven not by individual methanogens alone, but by complex, depth-resolved interactions among microbial guilds. Notably, the strongest predictors of soil porewater CH₄ concentrations were often upstream carbon-oxidizing organisms such as iron reducers (*Zikibacteria, Eisenbacteria, Geomobilibacter*) and fermenters (*Aminicenantales*) that likely support diverse functional methanogens (*Methanoregula, Methanothrix, Methanomethylicus*). These findings underscore the need for higher-throughput tools to resolve microbial interactions in the context of ecosystem-scale methane dynamics.

## Discussion

### Depth-structured microbiomes and redundancy shape wetland stability

Our study reveals that depth is the dominant axis structuring microbial composition and transcriptional activity in freshwater wetland soils. Across spatial and temporal gradients, microbial communities and biogeochemical functions were vertically stratified, with functional guilds consistently aligned with depth-defined redox and geochemical profiles. Traits such as methanogenesis, iron reduction, and fermentation were expressed by phylogenetically diverse organisms within the same depth layers, indicating high transcriptional redundancy. This redundancy, both within and across lineages, likely buffers methane-cycling functions against environmental fluctuations and contributes to the long-term stability of CH₄ production. While previous inferences of functional redundancy were drawn from alpha diversity or genome-level data (32), our findings demonstrate this redundancy at the level of gene expression and functional traits, offering mechanistic evidence of activity-based ecosystem resilience.

Contrary to the microbial seed bank hypothesis (33), we observed widespread transcriptional activity across archaeal and bacterial genomes. Specifically, 78% of the 2,502 genomes recruited transcripts, with 60% of these MAGs expressing sufficient genome content to permit metabolic assignments to 14 traits associated with wetland biogeochemistry. Moreover, the average MAG mean transcriptional occupancy was 59%, signifying active members were maintained across samples that span diverse wetland gradients. These patterns suggest that, rather than lying dormant, a large fraction of the wetland microbiome is metabolically active and responsive across depth and time. While functional redundancy may provide ecological resilience, it also complicates methane mitigation. In highly redundant systems, inhibiting one lineage or process may simply lead to functional compensation by others, a microbial game of ‘whack-a-mole’. As such, successful mitigation strategies must move beyond single-target interventions and account for the depth-resolved, community-level architecture of soil microbiomes.

### Expanding the methane paradigm: beyond classical substrates and lineages

This study also expands the framework of methane cycling by identifying alternative metabolic routes and underappreciated contributors. Although acetoclastic and hydrogenotrophic methanogenesis dominate conventional models (34), nearly a quarter of transcriptionally active methanogens in our study expressed genes for methylated compound utilization. These findings suggest that CH₄ production is not just fueled by acetate or hydrogen but also by methylated nitrogen, sulfur, and oxygen-containing substrates. Moreover, methane concentrations in porewater were more accurately predicted by co-expression networks involving iron reducers, obligate fermenters, methane oxidizers, and microaerophilic heterotrophs than by methanogens alone. These data reveal methane as an emergent property of integrated, depth-specific microbial consortia. In surface soils, aerobic methanotrophs likely attenuate CH₄ as its produced (based on the strong spatial co-expression), while deeper modules are dominated by organisms facilitating substrate production and syntrophic hydrogen production (**Fig. 6**).

These findings reinforce the need for methane mitigation strategies that are not only taxon-specific but also spatially and functionally informed. Interventions that ignore the trophic structure and spatial organization of microbial communities risk being ineffective or short-lived. For example, in other methanogenic systems such as the rumen (35, 36), mitigation approaches increasingly target upstream methanogen metabolic partners, manipulating redox conditions, or rerouting electron flow to non-methanogenic microbes. Similarly, recent work has shown that catechin, a naturally occurring plant metabolite found in wetland soil microbiomes, can suppress methane production by up to 84% by redirecting hydrogen flux away from methanogens (37). Especially in complex environments like saturated soils, precision strategies that consider microbial interdependencies, metabolic plasticity, and depth-resolved activity are likely to offer more robust and scalable outcomes for methane mitigation.

### Implications for redox-informed Earth system models

Our genome-resolved transcriptional profiles reveal seasonal and multi-year stability in methane-cycling activity across soil depths, despite substantial hydrological and redox perturbations. While we previously reported elevated CH₄ production under oxygenated conditions at this site in 2015 (15), new flooding data reinforce that methanogen transcription is largely unresponsive to shifts in soil oxygen concentrations. These findings add to a growing body of evidence challenging the long-standing paradigm that oxygen is the primary regulator of wetland CH₄ production (32–35). In our data, methanogen transcripts persisted in surface soils with detectable oxygen and showed no significant change following a 5.3-foot flood-induced redox shift, while aerobic methanotrophs remained active even under anoxic conditions. This decoupling suggests that microbial populations may be insulated within anoxic microsites, such as aggregates or biofilms, buffering them from bulk redox conditions. These results expose the limitations of using coarse redox proxies like dissolved oxygen in Earth system models and underscore the need for finer-scale, spatially explicit observations to resolve methane-cycling heterogeneity in wetlands.

Current biogeochemical models also largely neglect methylotrophic methanogenesis, despite mounting evidence that transcriptionally active methanogens frequently utilize methylated nitrogen-, sulfur-, and oxygen-containing compounds (16, 27, 38–40). These substrates, which we show are abundant in situ, may represent a substantial and previously under-accounted source of CH₄ emissions. To incorporate these pathways into predictive frameworks, targeted cultivation of methyl-reducing methanogens is needed (39, 41), particularly from wetland environments where cultured representatives remain scarce. Such efforts are essential for generating key physiological parameters (e.g., uptake kinetics, growth optima, yields), paralleling existing knowledge for acetate- and hydrogen-utilizing methanogens.

In parallel, stable isotope-based approaches will be vital for tracing substrate fate and quantifying the relative contribution of methylotrophy to CH₄ emissions. To help focus these efforts, our transcriptomic analysis distilled 85 methanogens into a “most wanted” list of four genera (*Methanoregula*, *Methanothrix*, *Methanomethylicus*, and Methanospirillaceae UBA9949) based on their transcriptional activity and predictive value for CH₄ concentrations (**Fig. 1**, **Fig. 6**).

Supporting this prioritization, a recent cross-site comparison across nine temperate wetlands (29), including Old Woman Creek, found *Methanoregula* and *Methanothrix* to be core taxa, with *Methanoregula* consistently emerging as a network hub strongly predictive of CH₄ flux. As methane remains a powerful lever for near-term climate mitigation (42), advancing the biochemical and ecological understanding of these keystone methanogens, and the environmental controls shaping their activity, is paramount for improving process-based models and enabling targeted interventions across wetland ecosystems.

Accurately predicting methane flux in wetlands requires a shift toward models that capture the functional and spatial organization of microbial communities. Recent advances, such as the genome-to-ecosystem framework, have shown that incorporating genome-inferred biogeochemical traits weighted by community composition can markedly improve methane predictions in peatlands, reducing model bias by more than 50% (43). While impactful, these models rely solely on genomic potential, which our findings suggest may fail to capture the biogeochemical pathways actively driving emissions. Our work builds on and extends this approach by integrating metatranscriptomic data to define trophic modules that reveal depth-stratified networks of carbon oxidizers, syntrophic cross-feeders, terminal methanogens, and methane oxidizers. By identifying the most active and predictive processes, our findings provide a foundation for developing in situ diagnostics that move beyond multi-omic sampling toward cost-effective proxies of microbial function. Embedding activity-based functional traits into Earth system models will be critical for improving methane predictions and informing scalable, precision-guided interventions across diverse wetland ecosystems.

## Conclusion

In this study, we employed a comprehensive, multi-omics survey of wetland soils to generate an unprecedented view of the structural and functional microbiome attributes that underpin biogeochemical cycling. This approach helps move soil microbiome research beyond the historical “black box” of environmental inputs and outputs, revealing how microbial community composition and gene expression respond to depth and geochemical gradients. To interpret this complexity, we developed gene-based rules to classify microbial energy metabolisms and carbon trophic guilds, enabling us to resolve emergent properties of methane-cycling consortia. This framework uncovered previously unrecognized roles for archaeal lineages in carbon decomposition, revealed the substrate versatility and redox resilience of methanogens, and demonstrated stable, depth-stratified co-expression patterns predictive of soil CH₄ concentrations. Together, these insights provide a mechanistic foundation for embedding microbial activity into Earth system models and for guiding microbiome-informed strategies to mitigate methane emissions from wetlands.

## Data Availability

The metagenomic reads, metatranscriptomic reads, bacterial and archaeal MAGs, and 16S rRNA gene sequencing reads reported here have been deposited in the National Center for Biotechnology Information (**Supplementary Table S1**). To facilitate data accessibility, the 2,502 MAGs contained in this dataset, here identified as MUCC (Multi-omics to understand Climate Change) v.1 are publicly available as a KBase data collection, offering genomes and annotations for use in an accessible cyberinfrastructure powered platform with the intent to advance wetland science broadly (https://narrative.kbase.us/narrative/147022). Flux and meteorological observations for OWC are available through Ameriflux, Site ID US-OWC (19). Porewater methane concentrations and chamber fluxes are available through ESS-DiVE (18) and (44). Data for nutrient and carbon sequestration and soil accretion rates of OWC is available through ESS-DiVE (45). Additional meteorological, hydrological and water quality data for OWC is available through the NERR data system (https://cdmo.baruch.sc.edu/). LC-MS data is available through MassIVE under accession MSV000093935. Additionally, all MAGs, and derived gene calls, annotations, phylogenetic trees, and metatranscriptome abundance profiles are available on Zenodo DOI 10.5281/zenodo.8194032.

## Notes

### Competing Interest Statement

The authors have declared no competing interest.

### Summary of Updates

Figures and text have been updated along with corresponding methods.

